# Cas9-induced large deletions and small indels are controlled in a convergent fashion

**DOI:** 10.1101/2020.08.05.216739

**Authors:** Michael Kosicki, Felicity Allen, Allan Bradley

**Affiliations:** Lawrence Berkeley National Lab, Berkeley, US; Wellcome Sanger Institute, Hinxton, UK; Department of Medicine, University of Cambridge, Cambridge, UK

## Abstract

Repair of Cas9-induced double-stranded breaks results primarily in formation of small indels, but can also cause potentially harmful large deletions. While mechanisms leading to the creation of small indels are relatively well understood, very little is known about the origins of large deletions. Using a novel library of clonal mouse embryonic stem cells *bona fide* deficient for 32 DNA repair genes, we have shown that large deletion frequency increases in cells impaired for non-homologous end joining and decreases in cells deficient for the central resection gene *Nbn* and the microhomology-mediated end joining gene *Polq*. Across deficient clones, increase in large deletion frequency was closely correlated with the increase in the extent of microhomology and the size of small indels, implying a continuity of repair processes across different genomic scales. Furthermore, by targeting diverse genomic sites, we identified examples of repair processes that were highly locus-specific, discovering a novel role for exonuclease Trex1. Finally, we present evidence that indel sizes increase with the overall efficiency of Cas9 mutagenesis. These findings may have impact on both basic research and clinical use of CRISPR-Cas9, in particular in conjunction with repair pathway modulation.

## Introduction

The goal of genome engineering is the introduction of particular genotype (or genotypes) to the exclusion of others. A programmable nuclease Cas9 is currently the primary tool of genome engineering in clinical and basic research context. Resolution of the double-stranded break (DSB) induced by Cas9 at a location determined by a guide RNA (gRNA) is the principal cause of Cas9 mutagenesis and templated editing. Since the specific outcome depends primarily on the relative activity of different DNA repair pathways, understanding of their function in genome engineering is crucial.

Cas9 mutagenesis is primarily the result of non-homologous end joining (NHEJ) and microhomology-mediate end joining (MMEJ) repair. NHEJ is initiated by Ku70/Ku80 complex binding to the ends of the break, protecting it from degradation. A cascade of events involving, among others, DNA-PKcs, 53BP1, Xlf, Xrcc4 and Lig4, leads to either a perfect repair or a small indel (<10 bp). Resection of the break by the Mre11-Rad50-Nbs1 (MRN) complex, promoted by Ctip and Brca1, prevents NHEJ. Resected DNA can be repaired through microhomology-mediate end joining (MMEJ), which involves Parp1, Polϑ and ligases Lig1 and Lig3, resulting in larger indels (1).

A frequency spectrum of indels resulting from NHEJ or MMEJ repair in a population of cells, the “indel profile”, is specific to local DNA sequence of the target and generally stable across tested cell lines (2–4). These indels, usually smaller than 50 bp, can be predicted from DNA sequence with high precision (5–8). In particular, frequent occurrence of 1 bp insertions templated from around the cut site has been linked to Cas9-induced DSBs with 1 nt 5’ overhang (9, 10). The size of indels is typically increased by NHEJ inhibition, while inhibition of the core MMEJ proteins, such as Parp1 and Polϑ, decreases them (3, 11–15). While predictable and partially malleable, Cas9 mutagenesis often does not lead to the desired genome engineering outcomes.

Cas9 templated editing, which hijacks the homologous recombination (HR) pathway, can lead to well-defined outcomes. If the cell is in S/G2 phase of its cell cycle and if the MRN-initiated resection proceeds further, the DSB can be repaired by HR using either sister chromatid (resulting in perfect repair) or an exogenously provided template. This process involves, among others, Brca2 and Rad51 (16). Templated repair using Cas9 is normally harder to achieve than mutagenesis and therefore a number of studies focused on increasing its frequency. In addition to optimization of transfection conditions and the template itself (e.g. 17, 18), inhibition of NHEJ proteins by pharmacological means is one of the preferred methods (e.g. 19–22). At least one company plans to test these inhibitors in the context of clinical genome engineering (23). Some of the alternative strategies include modulation of the cell cycle and of the HR pathway itself (24–28).

While a lot of literature has focused on small indels and templated repair, Cas9 complexed with a single gRNA can also induce large deletions at least kilobases in size and complex lesions, such as translocations, large insertion and noncontiguous lesions at significant frequencies (29–32). These effects were also noted in conjunction with templated editing in mice (33–36). Extensive loss-of-heterozygosity by gene conversion and megabase-long deletions were also observed (37–40). Such outcomes could be pathogenic, and may be hard to genotype. Methods which do not require DSB to introduce templated edits, such as base editing and prime editing, were developed in part to avoid such consequences. Nonetheless, these tools likely introduce DSBs occasionally, as evidenced by creation of indels, making it likely they can also introduce large deletions (41, 42). Furthermore, it is not well understood, which DNA repair pathways control their creation.

To avoid large deletions and complex lesion, we need to know which repair mechanisms lead to their creation. To study this issue, we have built a library of mouse embryonic stem cells *bona fide* deficient for 32 DNA repair genes, derived from a single clone constitutively expressing Cas9. Using this library, we discovered that NHEJ genes prevent large deletions, while the resection gene *Nbn* and the MMEJ gene *Polq* are necessary for their creation. We also find a strong correlation between the frequency of large deletions and size or microhomology usage of small indels, across a range of deficient clones. This implies that small indels and large deletions are controlled convergently. By targeting multiple genomic sites, we observed some gene deficiencies have highly locus-specific functions and discovered a new role for exonuclease Trex1. Finally, we have shown that highly efficient Cas9 editing leads to more MMEJ outcomes.

## Methods

### Generation of the Cas9+ embryonic stem cell clone

An EF1a-Cas9-T2A-blastR transgene in a pKLV backbone (2,43) was introduced by lentiviral transduction into a highly heterozygous CB9 mouse embryonic stem cell line, derived from a cross between CAST and C57BL/6 strains (44). Low titre of the virus was used to achieve low copy number. Blasticidin selected, single cell cloned colonies were isolated and tested for Cas9 efficiency using a flow cytometric assay with self-targeting BFP-GFP-anti-GFP construct (2) or a gRNA against *Cd9* gene (30). The most efficient and homogenous clone “CBA9” was picked for library creation.

All ES cells used in this study were propagated on SNL-blast feeder cells resistant to neomycin and blasticidin or SNL-HBP feeders resistant to neomycin, blasticidin, hygromycin and puromycin. SNL-HBP were created for this purpose by stable transposition of a low passage SNL cell line with a *PiggyBac* PGK-hygroR-blastR-puroR construct using hyperactive *PiggyBac* transposase (45) and selecting a multi-resistant pool of cells. Mouse ES cells grown on SNL-HBP feeders were found to have no morphological abnormalities compared to those grown on SNL feeder cells (data not shown).

### Generation of the DNA damage repair deficient library

*PiggyBac* transposons expressing a hygromycin resistance gene and gRNAs against DNA damage repair genes (Table S1, control, knock-out and knock-out-fail gRNAs) were introduced into CBA9 Cas9+ cells in an arrayed format using lipofection. Cells were selected for stable integration using 140 μg/ml hygromycin and single cell cloned. gRNA-targeted loci were amplified using barcoded primers (Table S1) and sequenced using MiSeq. Mutagenic alleles were called using CRISPResso2 (46) and manual curation of reads aligned using STAR (47). The latter method yielded additional large deletion alleles (>50 bp) missed by CRISPResso2.

Based on the recovered genotype, clones were classified as “perfect”, “in-frame” or “good”. Clones whose all detectable alleles were frame-disrupting and, where applicable, could be assigned to a strain (one BL6 and one CAST allele), were deemed “perfect”. These are very likely to have lost target gene function. Clones containing any alleles likely to be functional (frame-preserving insertion or deletions smaller than 30 bp) were considered “in-frame”, unlikely to have lost gene function, unless a critical protein domain was affected. Other clones, including those with more than two alleles, with one allele at loci without strain specific SNPs (potentially homozygous, or harboring a complex lesion undetectable by short-range PCR), with any in-frame deletions 30 bp or larger (likely deleterious) or with alleles that could not be assigned to a strain at a heterozygous locus (because the lesion erased the distinguishing SNPs), were classified as “good”. Control clones were obtained using various non-targeting gRNAs and a gRNA targeting a safe harbor *Rosa26* locus. In total, 57 perfect, 18 good and 8 in-frame experimental clones, along with 12 controls, were included in the final library for a total of 95 clones. One well was intentionally left empty as a negative control for cell and DNA carry-over. The library contained clones *bona fide* deficient for 32 genes, with 1-4 clones for each gene (Table S2). Two of these genes, *Brca2* and *Xrcc1*, were only represented by “in-frame” clones. No clones were included for seven other targeted genes, which yielded no promising candidates (*Brca1, Exo1, Mre11a, Rif1, Rnaseh2a, Fen1* and *Mad2l2*).

### Flow cytometric assessment of mutagenesis efficiency and complex lesion frequency

Flow cytometric assays were conducted as previously described (30, 48). In short, *PiggyBac* transposons expressing one of the five experimental gRNAs (#15,#33, #48, #48U and #148) and a puromycin resistance gene were introduced into the library clones in an arrayed format using lipofection. Cells were selected for stable integration using 10 ng/μl puromycin. This strategy ensures a near complete mutagenesis. On day 7 and day 14 post-transfection, cells were stained using FLAER reagent (for PigA activity; Cedarlane) or Itga6-PE antibodies (#313612, Biolegend) and analyzed using Cytoflex flow cytometer (Beckman-Coulter). Six replicates were performed for day 7 and four for day 14.

Data was extracted, processed and visualized in R, using packages flowCore, flowWorkspace, openCyto and ggcyto (49–51). The same gating was used throughout, except in replicate 6 on day 7, in which cells had to be restained and gates adjusted to lower staining efficiency. A bacterial infection was detected in replicate 1 on day 14 - cells were processed as usual and data was retained. Raw percentages of staining-positive cells from each plate (that is, replicate, experimental gRNA, staining combination) were mean and standard deviation normalized and resulting raw z-scores were decomposed using PCA. Two first principal components captured 60% of variation and separated the samples by gRNA and time of sampling. The next two components separated two batches of replicates (Fig.S1A). These batches were initiated from different master plates and used different lots of some reagents. The second batch grew faster (data not shown) and, possibly as a consequence, experienced an overall increased level of mutagenesis (Fig.S1B). We only used data derived from the first two principal components for analysis, expressed as a z-score with relation to mean and standard deviation of the control samples (e.g. in Fig.1B). Raw frequencies of gene expression negative cells, raw z-scores and PCA-regressed z-scores are presented side-byside in Fig.S2.

**Fig. 1.**
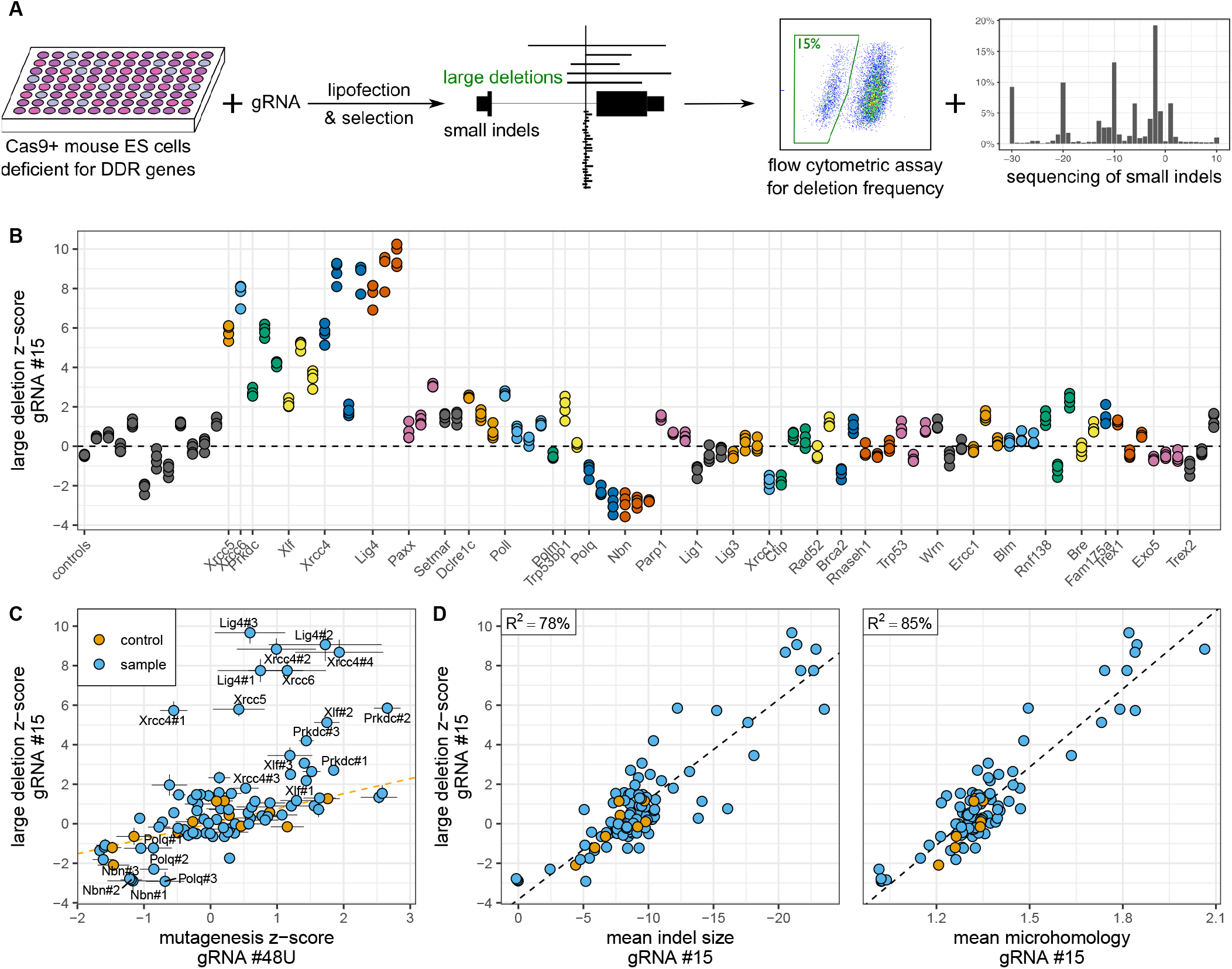
End joining pathways divergently control the frequency of large deletions caused by Cas9 in mouse ES cells. (**A**) Experimental design. Library of Cas9-positive clones *bona fide* deficient for DNA damage repair genes was transfected with individual gRNA-expressing constructs and selected for stable integration. Expression of target genes was measured by flow cytometry, revealing frequency of large deletions (using intronic gRNA #15) or overall mutagenesis (using gRNAs #48U, #48 and #148). Frequency of small indels was established by targeted sequencing of short-range PCR products. (**B**) Frequency of large deletions caused by Cas9 with intronic gRNA in DNA damage deficient clones, measured by flow cytometry, expressed as a regressed z-score (see Methods). Only the first clone in a series of clones deficient for the same gene is labeled on thex-axis. N=4 independent cell cultures. (**C**) Comparison of large deletion and mutagenesis indices (see Methods). Dashed line indicates best linear fit to control clones (in orange). Error bars are 2xSEM (N=4). (**D**) Correlation between large deletion z-score (measured by flow cytometry) and the size or microhomology extent of small indels (measured by targeted sequencing). Each dot represents an average readout of an individual clone (N=1-2 biologically independent cell cultures). Negative indel sizes indicate dominance of deletions.

### Analysis of indel profiles

Loci targeted with five experimental gRNAs in 95 clones in two biological replicates (day 14, replicates 5 and 6 in the flow cytometric assays) were amplified using barcoded primers (Table S1; amplicons of 244-283 bp) and sequenced using MiSeq. Demultiplexed reads were handled as described in Allen et al. (5). In brief, reads were transformed into indel signatures characterised by their size, type and location with respect to the cut site (3rd/4th nucleotide 5’ of the PAM). For example, ‘D10_L-13C2R0’ is a 10 bp deletion (‘D’), whose last unmodified nucleotides are thirteen to the left of the cut site and at the cut site. Two nucleotides of microhomology (‘M’) could map at either end of that interval. When an indel contains insertions (‘I’), microhomology indicates that this part of the insertion matches at either end of the interval, indicating a possible templated insertion.

Samples transfected using control gRNA #33 targeting GFP were found to contain negligible amounts of indels at #15, #48 and #148 sites and were discarded (data not shown). The following filters were applied to the remaining 760 samples. Indels that would result in loss of more than 150 bp were filtered out to avoid primer-dimers (0.6% read loss) and samples with fewer than 400 remaining reads were removed (18/760 samples lost; each gRNA-clone combination retained at least one sample). Symmetrized Kullback-Leibler divergence (“KL”) was calculated for all pairs of samples as described in Allen et al. (5). The resulting KL matrix was decomposed using multidimensional-scaling (MDS) for visualization and statistical testing. A bivariate normal distribution was fitted to controls using two first principal components of the MDS-decomposed KL matrix (Fig.S3A) and the associated p-value for each sample was calculated. An indel profile of a clone was considered significantly different from controls, if all of its replicates had a FDR-corrected p-value<0.01. Analysis and visualization were performed in R using ggplot2, ggrepel (https://ggrepel.slowkow.com/) and tidyverse group of packages, as well as colorblindr and cowplot (52, 53).

## Results

### Mouse embryonic stem cell DNA damage repair deficient library

Many DSB repair deficient cell lines have been used to study Cas9 engineering outcomes in the past. However, available lines have often been derived independently, from different mouse strains, tested using different Cas9 vectors and compared to only one or few control clones. This may introduce cryptic variability in such parameters as proliferation rate or Cas9 expression levels, which in turn may confound the effect of DNA repair deficiency. Likewise, recently developed pooled CRISPR drop-out screens are vulnerable to proliferation rate differences that are not related to tested phenotypes. Seeking to avoid these confounders, we built a mouse embryonic stem cell DNA damage repair deficient library, based on a single clone constitutively expressing Cas9. Mutations in 39 repair genes were introduced using Cas9 complexed with 81 gRNAs. We selected these genes to broadly cover the main DSB repair pathways, NHEJ, MMEJ and HR. We also included a number of exonucleases, expecting some of them might have a role in generating large deletions.

Approximately 800 clones (incl. controls) were created in a single experiment and passaged together, minimizing differences due to handling and reagents batches. Clones with no detectable wild-type allele and no frame-preserving indels at the target site were incorporated into the library (with few exceptions, see Methods and Table S2). As expected, attempts to mutate a number of HR genes resulted in extensive lethality and an increased number of in-frame indels (data not shown, see Methods for details). For some of these genes, we have incorporated clones with large in-frame deletions (more than 10 bp), expecting them to be hypomorphic. In total, we have selected 83 individual clones, with mutations in 32 repair genes. The library also included 12 control clones transfected with non-targeting gRNAs or a gRNA targeting safe harbor *Rosa26* locus (Table S2).

### Large deletions are prevented by NHEJ and promoted by *Nbn* and *Polq*

Large on-target deletions and complex lesions are a significant and potentially pathogenic outcome of Cas9 mutagenesis, but DNA repair pathways contributing to these outcomes are unknown. To map out these pathways, we have applied a previously developed flow cytometric assay, which detects such lesions (30), to the newly created arrayed library of clonal mouse embryonic stem cell clones *bona fide* deficient for DNA damage repair genes. We transfected each clone with a gRNA against the intron of the *PigA* gene and measured the frequency of cells that have lost PigA expression (Fig.1A). As shown before, small indels at this site do not affect PigA expression, and the cells that have lost gene expression harbor large deletions (>260 bp) overlapping the nearest exon or, much more rarely, other complex lesions which explain expression loss (translocations, non-contiguous lesions, insertions containing polyadenylation signals). We will refer to these events collectively as “large deletions”. The fact that in male ES cells there is only one copy of *PigA*, which is located on chromosome X, makes the assay highly sensitive.

We observed a substantial increase in large deletion frequency in clones deficient for the core NHEJ-factors, in particular *Xrcc4, Lig4, Xrcc5* (Ku80 protein), *Xrcc6* (Ku70), *Prkdc* (DNA-PKcs) and *Xlf* (Fig.1B). Mutations in other NHEJ genes, such as *Paxx, Setmar, Dclre1c* (Artemis) and *Poll* did not substantially influence the results, consistent with a minor role they play in this pathway. Conversely, lower frequency of deletions was found in clones mutated at the *Nbn* locus (Nbs1 protein), which is involved in initial resection of DSB leading to MMEJ and HR pathways. Similarly, deletions were less common in clones deficient for *Polq* (Polθ), a crucial component of MMEJ pathway. Raw frequency of large deletions spanned from almost 30% in *Xrcc4* and *Lig4* deficient clones to around 1% in *Polq* and *Nbn* deficient ones, compared to 12% in control clones. Since our assay primarily detects deletions spreading in a single direction from the cut site, the true frequency of these lesions is likely 1.5-2 times higher than measured (48).

To control for the expected variability in mutagenic efficiency between individual clones, we compared the deletion frequency with results obtained using exonic gRNA #48U, which tracks mutagenic efficiency. We chose this gRNA as reference, since mutagenesis using other exonic gRNAs was nearing saturation (#48 and #148; Fig.S2B, raw frequency). The effect on deletion frequency generally exceeded that on the overall mutagenesis level (Fig.1C). We conclude that large deletions are prevented by NHEJ repair and promoted by at least some part of MMEJ machinery.

### Small indels and large deletions are controlled by the same pathways

The effect of DSB repair pathways on large deletion frequency was qualitatively consistent with the previously described effect of these pathways on local indel profiles. A close quantitative correlation between the two would imply a common mechanism. To see if this is the case, we sequenced a 283 bp area around the cut site of the gRNA #15 we have used to assess large deletion frequency. We found a strong correlation between the average size of microhomology of the sequenced small indels and large deletion frequency as measured by flow cytometry (Pearson *R*^2^=85%, Fig.1D). We also found a moderate, inverse correlation with the average size of the small indels (*R*^2^ =78%, Fig.1D). A linear model using both measures was not significantly different from single-measure models (data not shown). These observations imply a strong commonality of repair mechanisms generating both types of lesions and suggest that sequencing of short-range PCR products could be developed as an alternative assay for large deletions.

### Core end joining genes influence indel profiles of multiple target sites

Screens of DNA damage repair processes often rely on a single locus reporter assay or on composite readouts based on random mutagenesis. However, *in vitro* biochemical studies show that DNA repair is often highly sequence specific. To distinguish between universal and specific repair processes, we sequenced mutagenized target sites of three gRNAs, each with a distinct indel profile in control clones (Fig.2A). In particular, gRNA #15 was characterized by preponderance of 1 bp insertions, gRNA #48 by diversity of small indels 1-5 bp in size, while gRNA #148 induced discretely sized deletions (2 bp, 10 bp, 20 bp). We speculate that these profiles reflect relative contribution of NHEJ and MMEJ repair at a given site.

**Fig. 2.**
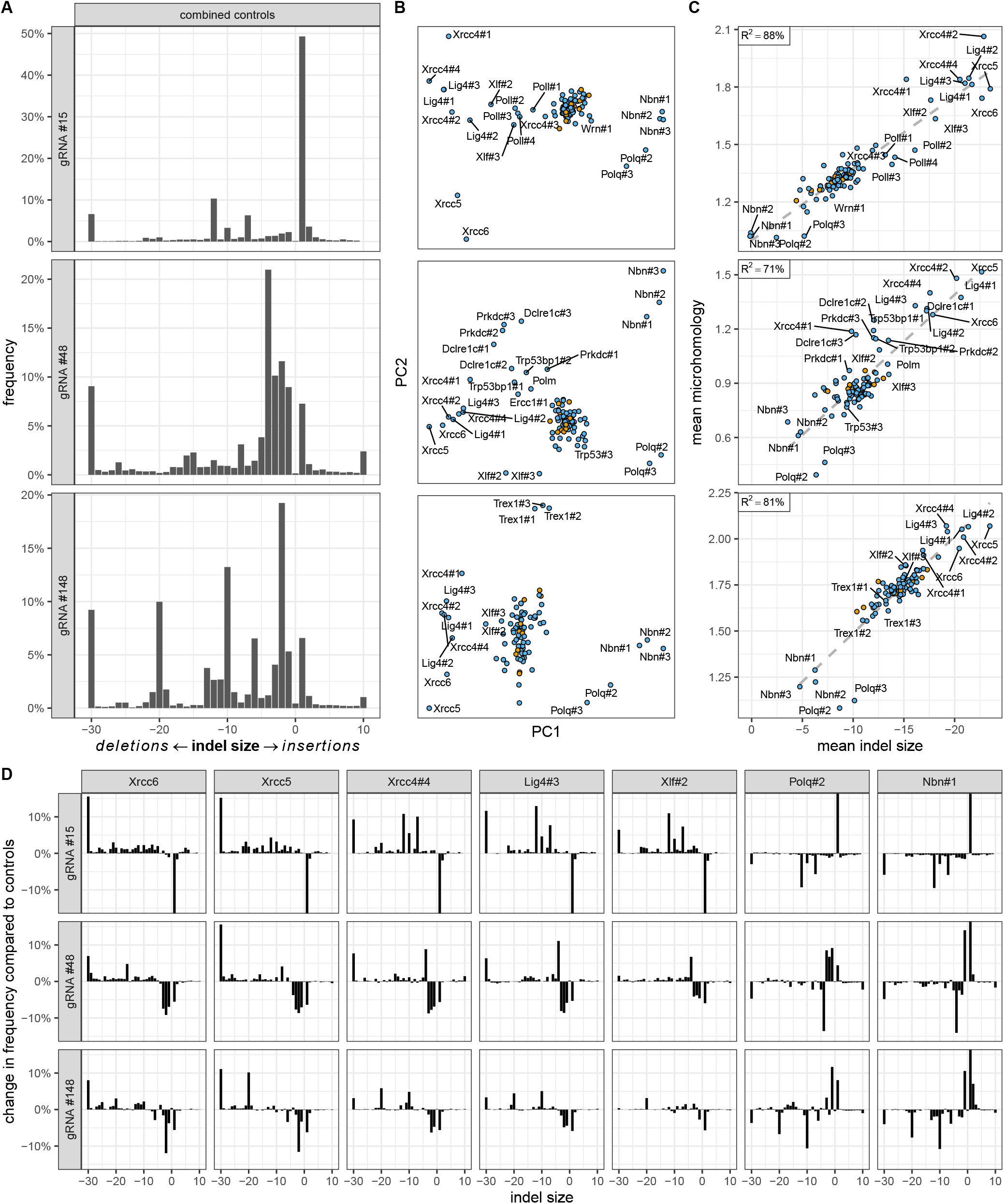
Core end joining genes influence indel profiles globally. (**A**) Indel profiles in combined 12 control samples. (**B**). Relationships between cell clones based on their indel profiles. Clones significantly different from controls (p<0.01) are labeled. The arrangement of non-significant clones is in Fig.S3C. (**C**) Correlation between mean indel size and microhomology. (**D**) Relative frequencies of indel sizes compared to controls in deficient clones with a significant impact on all three gRNAs. Indel profiles of other clones with significant impact are in Fig.S4. Y-axis is truncated at −15% and +15%. In panels A and D, indel frequencies are aggregated by combined size. Negative numbers represent deletions and positive ones represent insertions. The leftmost and rightmost bars (−30 and 10) combine all larger deletions and insertions, respectively. Biological replicates (N=2) were averaged for clarity. All rows in panels B and C relate to the same gRNAs as in panel A. In panels B and C, controls are in orange and samples are in blue.

To obtain an overview of relationship between deficient clones, we have calculated Kullback-Leibler divergence between each pair as described in Allen et al. (5) and transformed the resulting divergence matrix using multidimensional scaling (MDS), a non-linear dimensionality reduction technique similar to principal component analysis (PCA). We found biological replicates to cluster together, indicating good reproducibility (Fig.S3A). Furthermore, the majority of clones, including all controls, clustered at the centre of the plot. This indicated most mutants did not influence the indel profile substantially, consistent with the flow cytometry assay. As a further control, we compared the frequency of mutated reads and the frequency of cells which lost expression of the target gene in the flow cytometric assay and found them to match closely for exonic gRNAs #48 and #148 (Fig.S3C). As expected, these numbers did not match for the intronic gRNA #15, as in this case the two methods measure mutually exclusive outcomes: the frequency of small indels and the frequency of large deletions.

We asked which deficiencies exhibited similar effects regardless of the target site, and which other deficiencies they clustered with. Mutations in *Xrcc5* and *Xrcc6* genes, whose products form a functional heterodimer (Ku80-Ku70), had very similar, strong effects (Fig.2B and Fig.2D). Likewise, indel profiles of *Xrcc4* and *Lig4* mutants clustered together, consistent with the fact Xrcc4 forms a scaffold for Lig4. MMEJ-associated *Polq* and *Nbn* clustered away from NHEJ genes such as *Xrcc’s* 4,5 and 6 and *Lig4*. As shown previously, NHEJ-deficiencies increased the size of indels, while MMEJ-deficiencies decreased them, although specific role of *Nbn* in indel profile modulation has not been described previously. In general, genes acting earlier in their respective pathways (*Xrcc5*/*Xrcc6* and *Nbn*) had stronger phenotypes than the genes acting later (*Lig4*, Xrcc4 and *Polq*).

Resection exposes single-stranded DNA, which can participate in repair using microhomology. The extent of microhomology in an indel profile could thus be confounded by the extent of resection. Taking advantage of the wide range of repair outcomes in both control and deficient clones, we decided to investigate the relationship between the two. We found a striking correlation between the average indel size (proxy for resection) and microhomology size for all gRNAs (Pearson *R*^2^ between 71% and 88%, Fig.2C). On the average, we observed 1 bp more homology for 19-27 bp increase in indel size (depending on gRNA), with the caveat that we do not know if this relationship can be extrapolated beyond the observed intervals. We speculate that this perspective may allow assessment of the relative contribution of deficient genes to resection and microhomology repair, respectively. In particular, we think it is likely that clones close to the regression line (*Trex1, Nbn, Lig4, Xrcc5* and *Xrcc6*) mainly control the extent of resection, while “distal” clones (Polq, some of the significant *Xrcc4* clones, *Dclre1c, Prkdc* and *Trp53bp1*) also control the extent of microhomology, at least in some genomic contexts. Consistently, Polϑ, the gene product of the most systematically “regression line-distal” gene, is known to actively generate homologous DNA at the DSB ends. We note that clone Ercc1#1 with gRNA #48 was excluded from this analysis as a strong outlier, with much larger mean deletion size than controls (−38 bp), without a proportional increase in microhomology usage (1.2 bp). This may have been a consequence of insufficient read depth (1400 mutated reads) and lack of replicate sample, so this results must be treated with caution.

### Specialized repair pathways affect indel profiles in a locus specific manner

Having focused on indel patterns that were common between the three gRNAs, we turned to gRNAs-specific effects. We found that *Poll* deficiency only had a significant effect on the profile of gRNA #15, *Trex1* on #148 and *Prkdc*, *Dclre1c*, *Trp53bp1*, *Polm* and *Ercc1* on #48 (Fig.2B, examples of differential indel profiles in Fig.3A). Clones *Wrn#1* and *Trp53#3* also had specific effect on gRNAs #15 and #48, respectively, but it was far weaker than that of other genes and did not replicate in other independently derived clones. We chose not to explore this further.

**Fig. 3.**
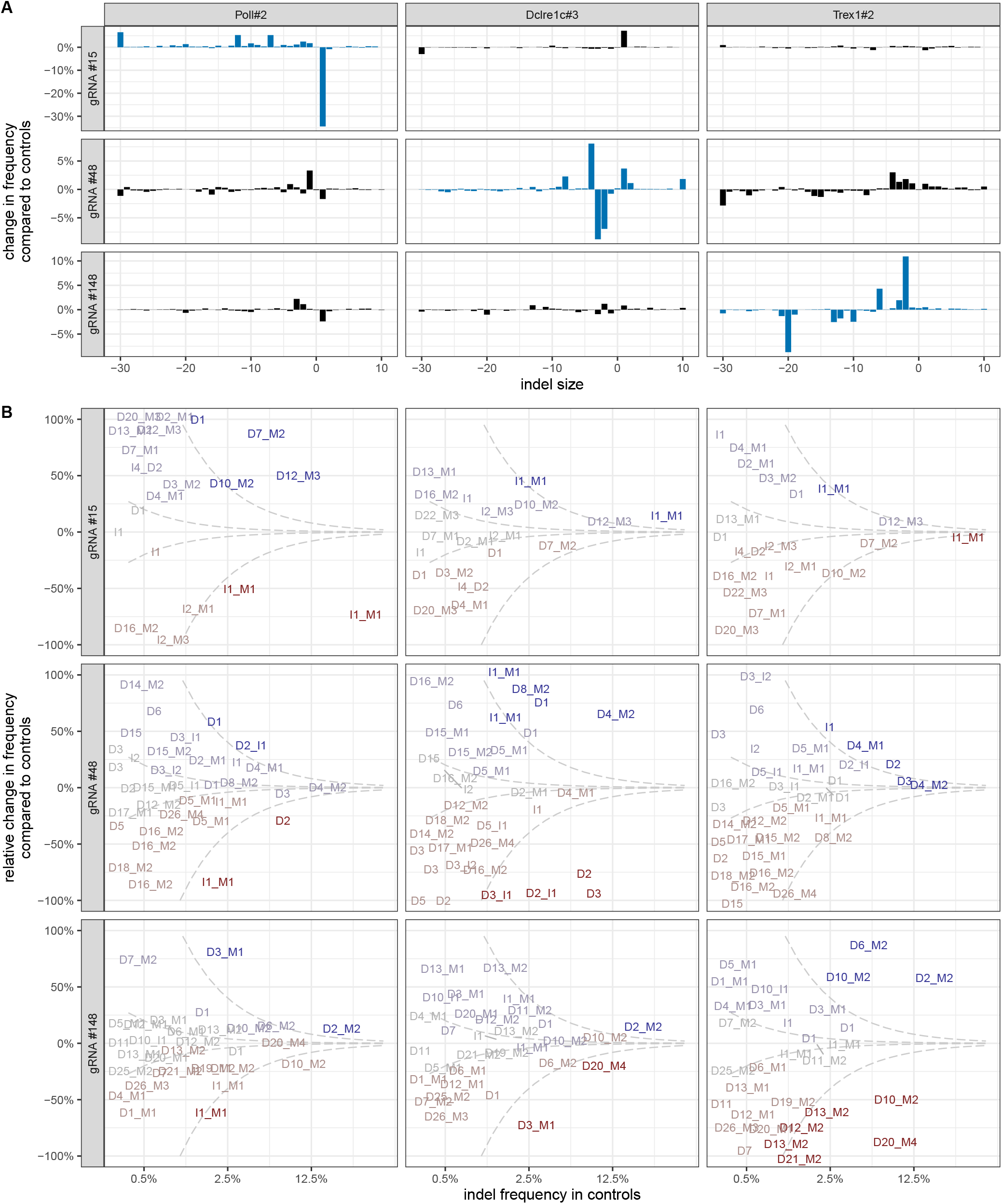
Deficiencies in specialized DNA damage repair genes influence indel profiles in a locus-specific manner. (**A**) Indel profile divergence between controls and selected clones. Blue bars highlight clone/gRNA combinations that were significantly affected (p<0.01). (**B**). Change in frequency of individual indels relative to controls. ‘D’ = deletion, ‘I’ = insertion, ‘M’ = microhomology (see Methods). X-axis indicates the frequency in control clones. Y-axis indicates relative change in indel frequency in a given clone relative to control clones. Complete loss of the indel is at −100%, while 100% indicates doubled frequency. The axis is truncated there for display clarity. Only indels present at 0.3% frequency or higher in control clones are shown. Dashed lines indicate absolute change of 0.1% and 1% respectively, color gradations highlight this change. Biological replicates (N=2) were averaged for clarity. All columns relate to the same gRNAs as in panel A.

We speculated this gRNA specificity is driven by the most prominent indels in each profile. By examining individual indel frequencies we confirmed that 1 bp, microhomology-associated insertions depleted by *Poll* deficiencies in profiles of all tested gRNAs, were most common in the profile of the significantly affected gRNA #15 (Fig.3B). Analogously, microhomology-containing 7-20 bp deletions prone to *Trex1* depletion were the most prominent outcomes of #148 mutagenesis. Finally, top indels depleted by *Prkdc*, *Dclre1c*, *Trp53bp1*, *Polm* and *Ercc1* were 2-5 bp deletions, commonly induced by gRNA #48 (Dclre1c example in Fig.3B). The effects of all deficiencies described here are consistent with the literature, except *Trex1*, whose function in DSB repair has not been described before.

To learn more about the effect of individual deficiencies, we examined indels that did not conform to the rules broadly laid out above. We found that gRNA #148 induced two different, prominent 10 bp deletions with 2 bp microhomology, whose frequency changed divergently in *Trex1* deficient clones. One of them, the only notable large indel to increase in frequency upon *Trex1* depletion, involved a G-homopolymer. Another divergent indel, a 4 bp deletion with 2 bp microhomology induced by gRNA #48, was promoted by deficiencies in *Prkdc*, *Dclre1c, Trp53bp1* and *Polm* (but not *Ercc1*), which otherwise decreased the frequency of 2-5 bp indels. We believe targeting additional loci to find more such apparently divergent outcomes could be useful to learn the rules governing DNA-sequence specific DSB-repair.

### Efficient mutagenesis leads to increase in size of small indels

Cas9 has a number of properties that make it likely to interfere with the DSB repair process. Among others, Cas9 can recut the DNA immediately after a perfect repair, may cut both sister chromatids simultaneously, stays bound to DNA after introducing the cut and might possess exonuclease activity. If Cas9 interferes with DSB repair in any of these ways, then manipulating its concentration or activity could result in changed indel profiles. To investigate this issue, we have challenged the library with a low efficiency gRNA #48U, whose target sequence is identical to #48. Unlike #48, #48U’s scaffold is expressed as two independent molecules, the crRNA (containing the target-matching sequence) and the tracrRNA. Significantly fewer control cells transfected with this weak gRNA lost PigA expression compared to the strong one (around 12% vs 80%, see Fig.S2). We speculate this is a consequence of reduced amount of “productive” gRNA.

To investigate the effect of mutagenic efficiency on repair outcomes, we first compared the results of the flow cytometric assay using gRNAs of different strengths. Samples transfected using the weak gRNA #48U clustered away not only from #15 samples, which track deletion frequency, but also from the combined cluster of strong exonic gRNAs #48 and #148 (Fig.4A). This difference was unlikely to be purely driven by the lower flow cytometry read-out with the weak gRNA, because the input for PCA-transformation was mean and standard deviation normalized, which should remove information about the relative magnitude of mutagenesis. Furthermore, #48U samples collected on day 14 posttransfection were further away from the #48 and #148 cluster than samples collected on day 7, which is contrary to the expectation of the observed principal components capturing the magnitude of mutagenic efficiency. We conclude that mutagenic efficiency qualitatively affects the results of the flow cytometry assay.

**Fig. 4.**
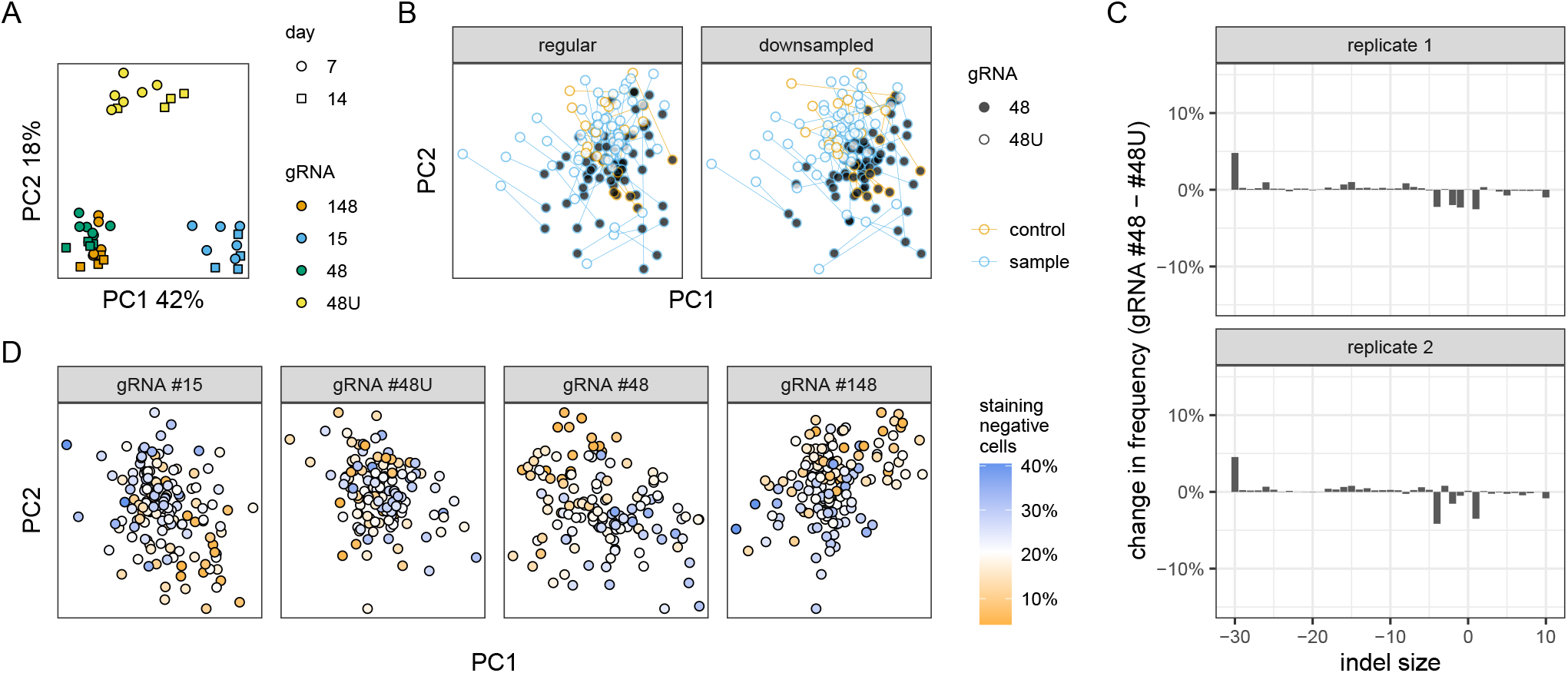
Efficiency of mutagenesis affects DNA repair outcomes. (**A**) Relationship between flow cytometry samples. gRNA #48U is a weaker version of #48. N=4-6 biologically independent replicates. (**B**). Relationship between clones based on their indel profiles, analogous to Fig.2B. Only non-significant clones are shown for clarity. Biological replicates (N=2) were averaged. (**C**) Difference in indel frequency between the regular gRNA #48 and its less active counterpart #48U. Same display conventions as in Fig.2. (**D**) Relationship between clones based on their indel profiles, analogous to Fig.2B. Each clone is colored by the frequency of mutagenesis assayed by flow cytometry on day 14 using gRNA #48U, a proxy for Cas9 activity. For clarity, only non-significant clones are depicted.

To test whether mutagenic efficiency affected small indel profiles as well, we compared the sequencing results of gRNAs #48 and #48U. The central cluster of controls and non-affected gene-deficient clones was clearly split between the two gRNAs (Fig.4B, left). Since mutated alleles were sequenced much more shallowly in #48U samples, which could potentially affect the results, we downsampled all read counts to the lowest common denominator (450 reads) and found that the effect persisted (Fig.4B, right). Indel profiles from combined control clones transfected with the strong gRNA had a higher frequency of larger deletions (5 bp deletions and larger), and correspondingly lower frequency of small indels, than clones transfected with the low efficiency gRNA (Fig.4C). This shift was reminiscent of one observed in NHEJ-deficient clones (such as Lig4 and Xrcc4) and could be interpreted as a relative increase in DNA resection and MMEJ-activity. The magnitude of the effect was small (no indel size changed in frequency by more than 5% percentage points), but reproducible between biological replicates.

There was considerable variability in the mutagenic efficiency among control clones in the flow cytometric assays (Fig.1B). We speculated that these differences in control and non-significant clones will correlate with differences between indel profiles. To ensure the highest dynamic range, we used day 14, gRNA #48U flow cytometry samples as a gauge, since in this sample only about 19% of the control cells are mutagenized. We found that mutagenic efficiency in this sample measured by flow cytometry correlated with divergence in indel profiles, as evidenced by the separation of clones in principal component space (Fig.4D).

## Discussion

We investigated the consequences of Cas9 mutagenesis in a panel of homogenous DNA damage repair deficient mouse embryonic stem cells. We found that the frequency of the complex lesions and large deletions (>260 bp) is increased by NHEJ deficiency and decreased by deficiencies in resection and MMEJ repair (Nbn and *Polq*). This is the first, systematic, quantitative assessment of this effect in a wide range of deficient clones, using an assay specifically designed to capture them. Large deletion frequency correlated with the increase in extent of microhomology and size of small indels. These result are consistent with the described functions of the identified genes and imply a continuity of underlying repair processes across large genomic distances. They also underscore the potential mutagenic danger of NHEJ inhibition, a common strategy for increasing the frequency of templated repair.

Our results also imply a strategy for decreasing the frequency of complex lesions, namely inhibition of MMEJ or resection, in particular by targeting *Nbn*. While global inhibition will decrease cellular viability and, in case of resection, genomewide repair fidelity (by preventing homologous recombination), a more targeted approach may be viable, like combining Cas9 enzyme with a resection-inhibiting moiety. Combining this strategy with prime editing could further reduce damage in rare cases when a DSB occurs. Another potential application of resection inhibition is to expedite the production of engineered cell clones by reducing the incidence of cryptic complex lesions (38). Moreover, repair outcomes in a resection-deficient context are much more predictable and more likely to lead to frame-disrupting 1-2 bp deletions and insertion. However, we note that the frame-shifting phenotype caused by 1 bp lesions may also be more likely to be rescued by genetic compensation (54).

A number of deficiencies in auxiliary repair genes had locus-selective effects. This observation is fully consistent with the well-described substrate-specificity of the repair genes involved, and the fact local target sequence shapes repair outcomes. Combining local nuclease-coupled manipulation of DNA repair machinery (e.g. 27) and indel profile prediction may be a viable strategy for obtaining the desired editing outcomes at a wide range of targets with minimal disruption to physiological DNA repair.

We were surprised to discover that *Trex1* deficiency had altered indel profiles. Trex1, discovered in 1969 and purified three decades later, has been studied extensively for its role in preventing autoimmunity caused by excess of ssDNA in the cytosol (55–59). Its *in vitro* exonuclease activity is fully compatible with a role in DSB repair but, to our knowledge, this is the first report linking the two. The underlying mechanism is unknown. Trex1 could potentially act upon DSB directly, for example during S phase, when it is involved in resolution of dicentric chromosomes (60, 61). Alternatively, the observed effect could be a secondary consequence of ssDNA accumulation, perhaps related to the increase in mutagenic repair upon transfection of non-homologous DNA reported by Richardson et al. (62).

We have shown that increased efficiency of Cas9-mediated mutagenesis correlated with MMEJ-like shift towards larger indels. We have noticed a similar effect before, when comparing different modes of Cas9 delivery (63). Multiple mutually non-exclusive causes for this are possible. More efficient Cas9 complex can recut the DNA after perfect repair sooner (potentially leading to chromatin state dependent repair modulation), cut both sister chromatids simultaneously more often (confounding HR repair), stay bound to DNA after introducing the cut longer on the average (64, perhaps interfering with the assembly of repair machinery or causing replication fork stalling or collapse) and exert its potential exonuclease activity more intensely (*in vitro*: 65, 66), than Cas9 of lower efficiency. Finally, the difference in observed profiles could in part be a temporary consequence of slower MMEJ repair dynamics. As long as new DSBs are being introduced, there is an excess of alleles under repair by the MMEJ pathway compared to the faster NHEJ pathway. Alleles in the process of being repaired cannot be amplified and are thus depleted from the observed indel profile. It is not trivial to figure out in which direction this process would push the indel profile, and what the magnitude of this effect is in our assay. An experiment using inducible gRNAs or inducible Cas9 of different strength would clarify this issue. However, we believe it is unlikely that the difference we observed is entirely due to this, as the effect persist, when mutagenesis is nearing saturation (Fig.S2, gRNAs #15, #48 and #148). Our results warrant further investigation and urge caution when using high concentrations of nucleases.

Despite deriving all our clones simultaneously from a pure, single cell cloned line, we observed a variability in mutagenic efficiency between control clones (e.g. 19-63% on day 7 with gRNA#48). The first round of subcloning likely removed most of the genetic variability, both genomic and related to individual lentiviral transductions (reverse transcription and APOBEC-mediated mutagenesis). Therefore, we speculate that differences in efficiency were due to a stochastic, mitotically heritable, epigenetic process acting on the Cas9 transgene, possibly position effect variegation. Since varying intensity of DSB introduction has influence on indel profile measurements, this might have precluded us from observing more subtle changes brought about by DNA repair deficiencies. Variation in mutagenic efficiency between Cas9 clones needs to be carefully consider as a potential confounder, when studying DNA damage repair.

Many genes in our library had no clones with statistically significant changes in indel profiles. Since their knock-out is only presumed based on the absence of small, frame-preserving indels, we cannot claim it as evidence of no function. We note that some clones *bona fide* deficient for core end joining genes, such as Xlf#1, Xrcc4#3 and Polq#1 exhibited a very mild or completely absent phenotype, while other clones with similar genotypes had very strong phenotypes. This, combined with the fact no significant effect was observed for genes with well-described functions in end joining, such as *Parp1, Lig1, Lig3, Ctip* and *Paxx*, implies genetic compensation might play a role.

## ACKNOWLEDGEMENTS

We would like to thank Lara Urban for help with running the PEER package, which in our case yielded the same results as PCA. Kärt Tomberg, Ilias Georgakopoulos-Soares, Ozdemirhan Serçin1 and Michał Barski offered useful comments on the manuscript. The bioRxiv version of the manuscript was typeset using a slightly modified Henriques Lab LATEX template. This work is funded by Wellcome Trust grant no. 098051.

## AUTHOR CONTRIBUTIONS

M.K. designed, executed, analyzed and interpreted the experiments and wrote the paper. A.B. supervised the project and contributed to writing of the manuscript. F.A. processed indel profile data and made useful comments on the manuscript.

---

**Fig. S1.**
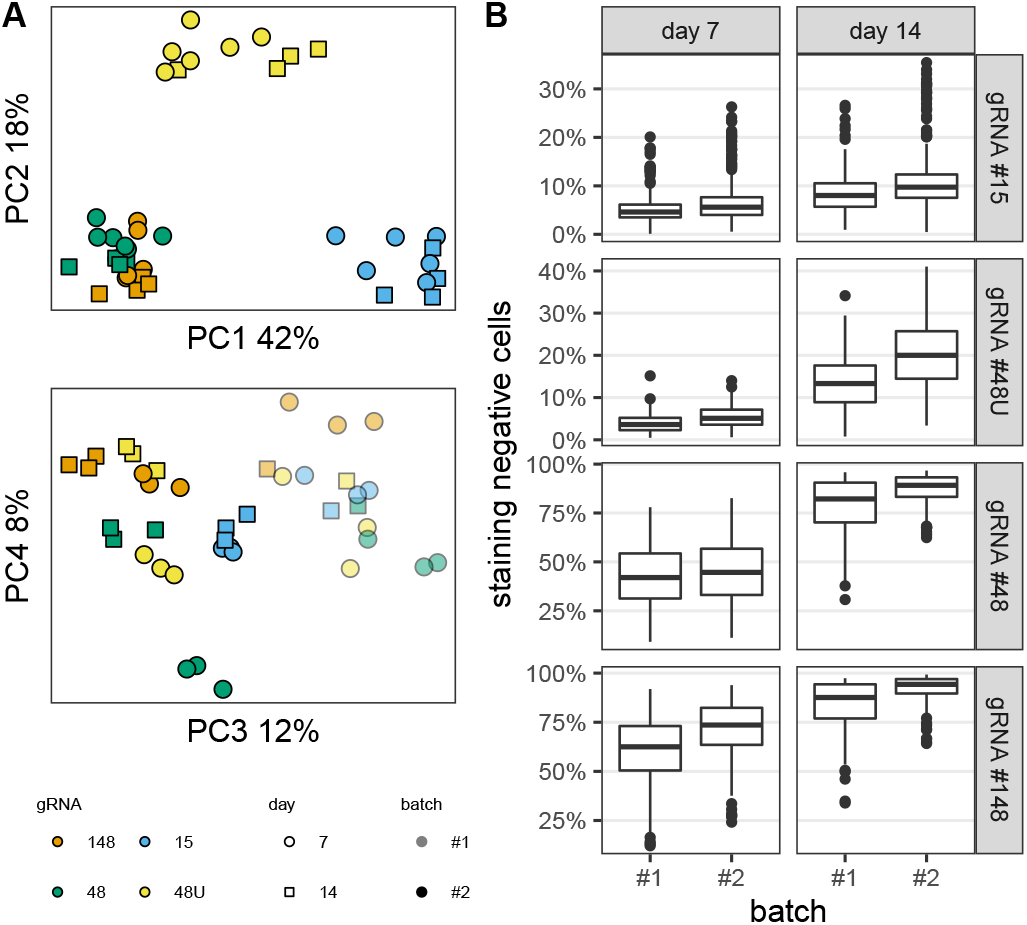
Batch effect in the gene expression assay. (**A**) Relationships between samples based on surface gene expression. (**B**) Frequency of cells negative for the expression of target genes, broken down by batch.

**Fig. S2.**
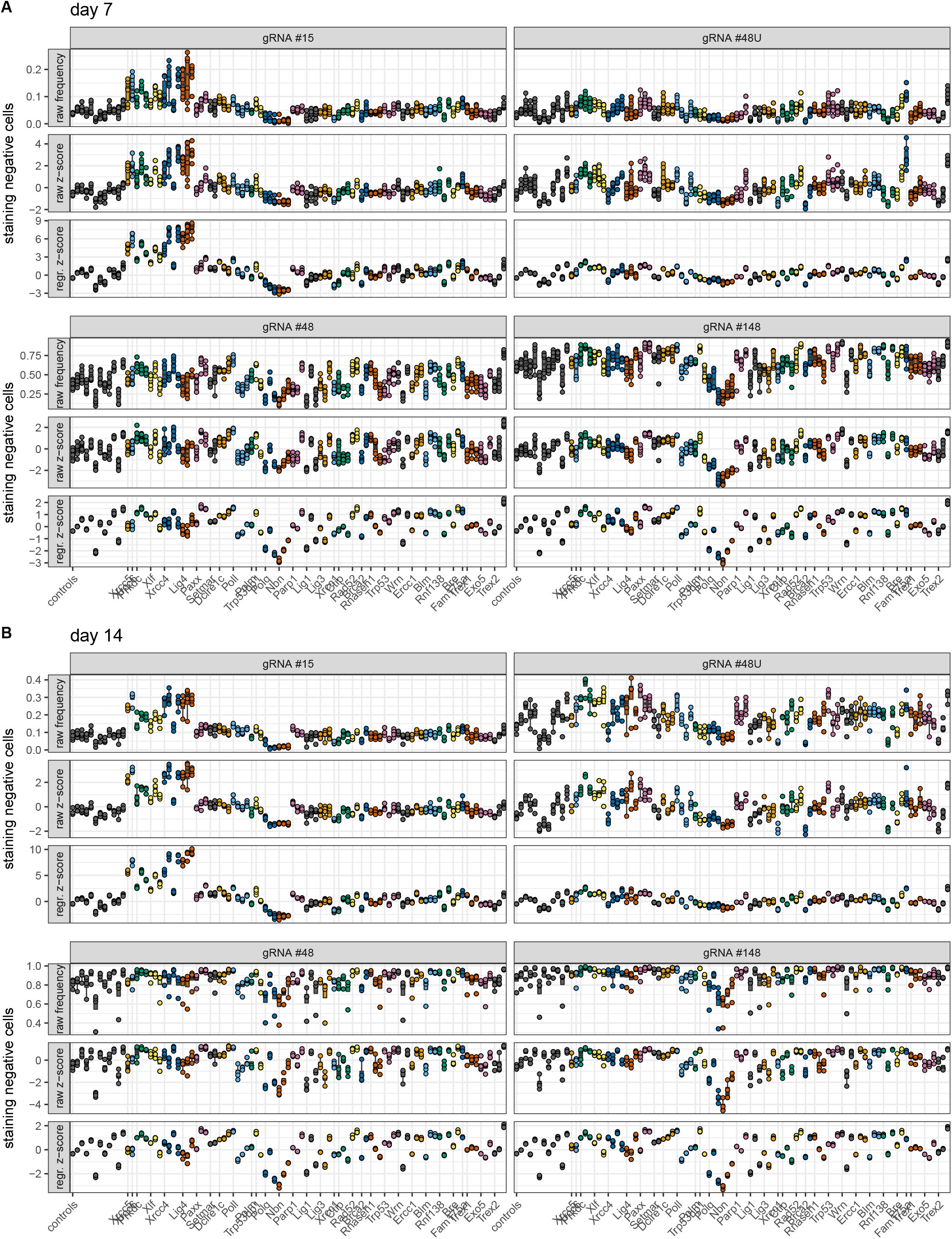
Gene expression negative cells - raw frequency, raw z-score and PCA-regressed z-score. (**A**) Day 7 post-transfection. (**B**). Day 14. Same display conventions as in Fig.1B.

**Fig. S3.**
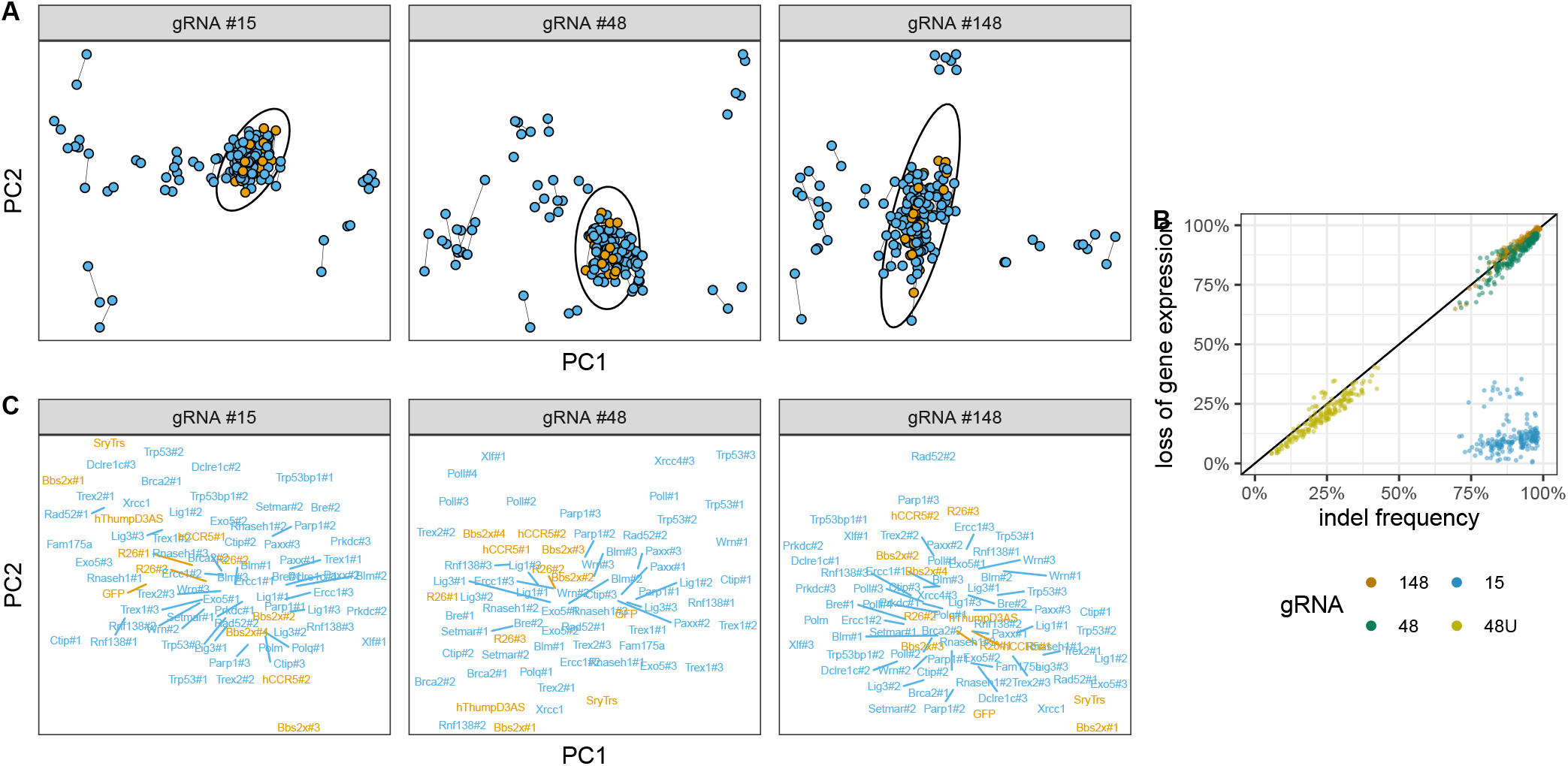
Assay reproducibility and relationships between non-significant clones. (**A**) Relationships between replicate measurements of indel profiles. Ellipse is 99% confidence interval of a bivariate normal distribution fitted to control clones (orange points). (**B**) Comparison of the frequency of mutated alleles as measured by targeted sequencing of short-range PCR products and flow cytometric measurement of protein expression. NB: only large deletions and complex rearrangements, and not small indels, result in loss of gene expression with intronic gRNA #15 (**C**) Relationships between clones not significantly different from controls, compare Fig.2B. Control clones are in orange. Biological replicates (N=2) were averaged.

**Fig. S4.**
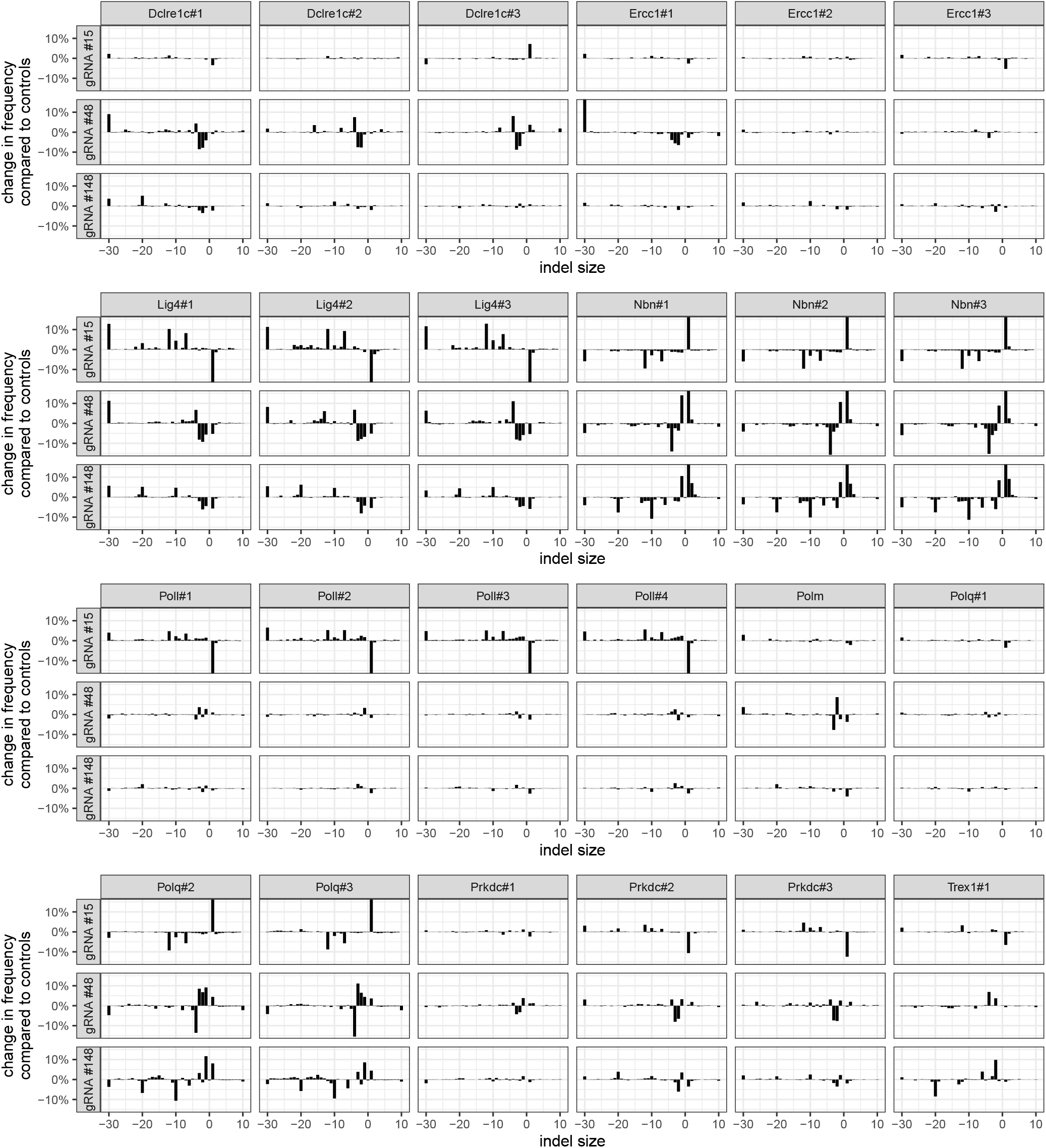
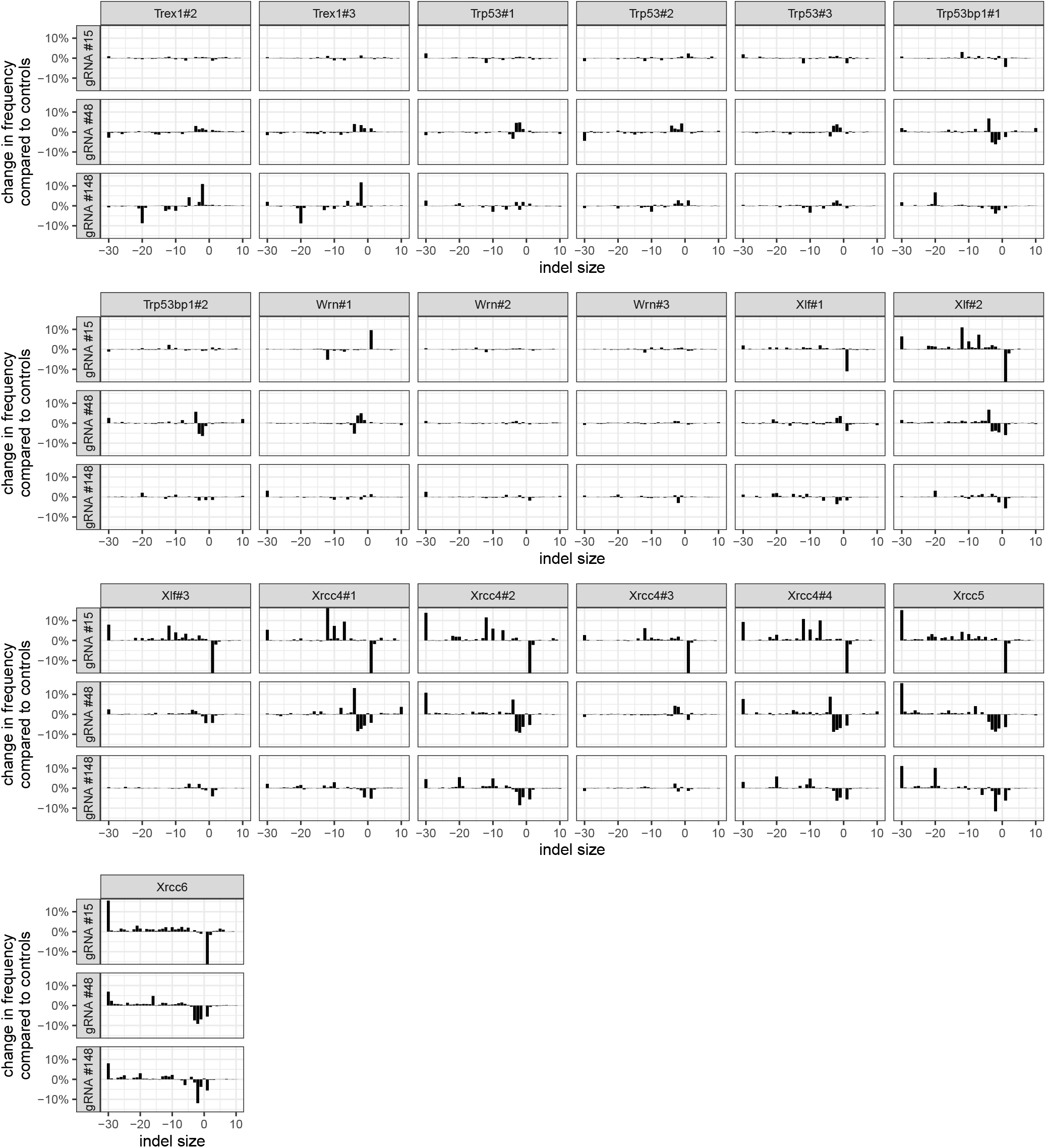
Indel profiles of genes with at least one significant clone. Y-axis is truncated at −15% and +15% for clarity. Figure spans two pages. Same display conventions as in Fig.2.

